## Supplementary figures and images for "Drug-induced increase in lysobisphosphatidic acid reduces the cholesterol overload in Niemann-Pick type C cells and mice"

### Supplemental Figures 1 to 10

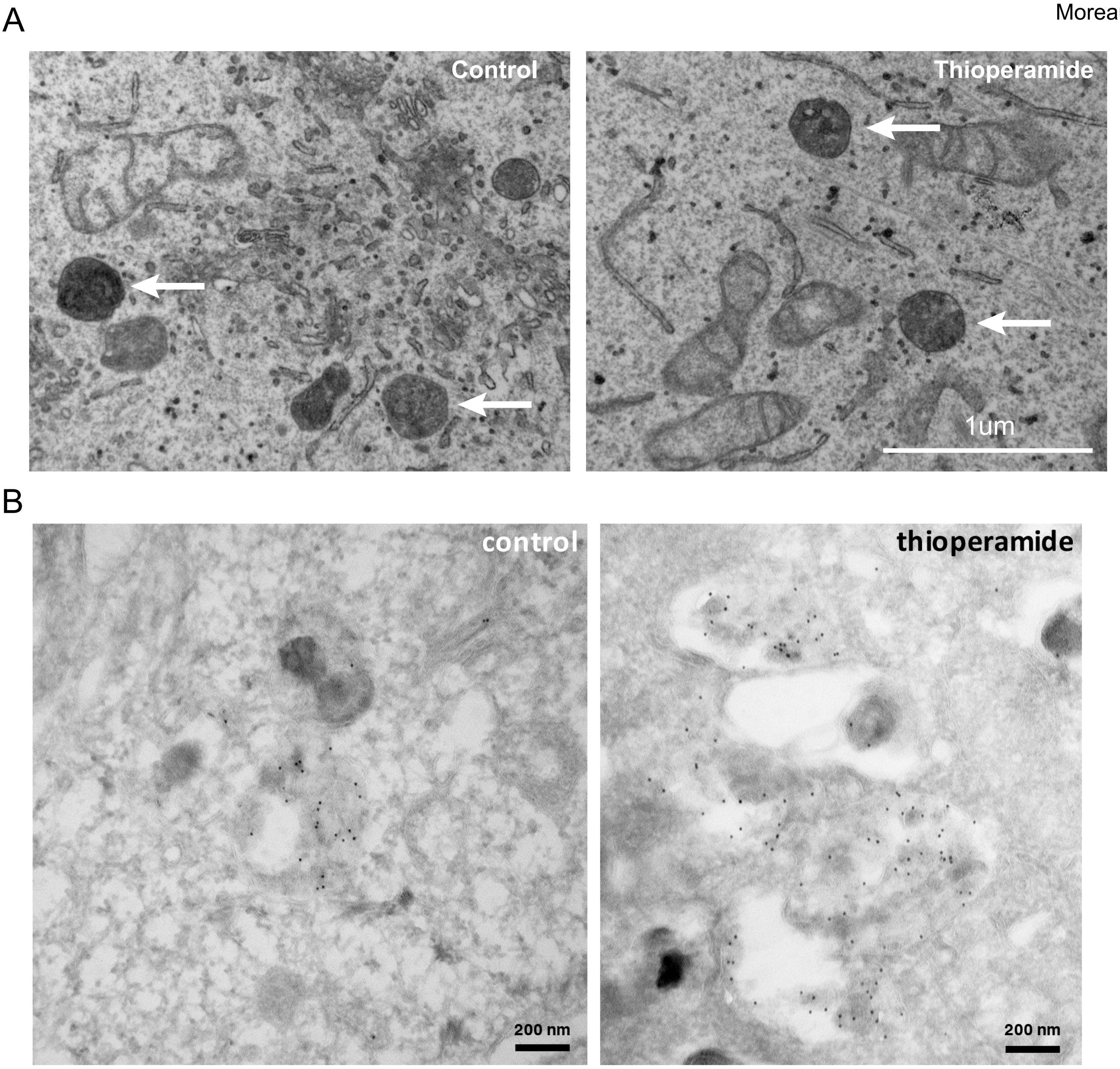

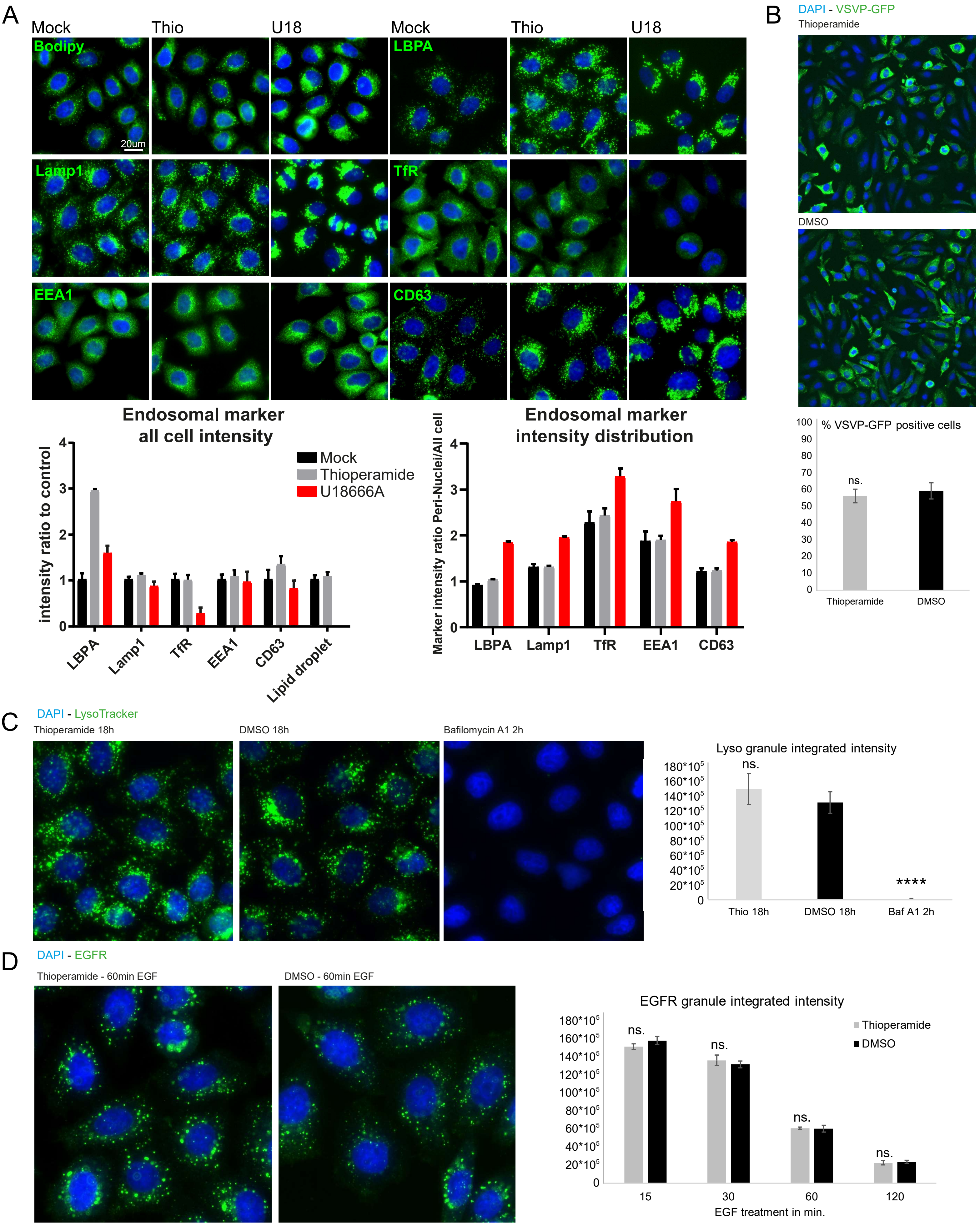

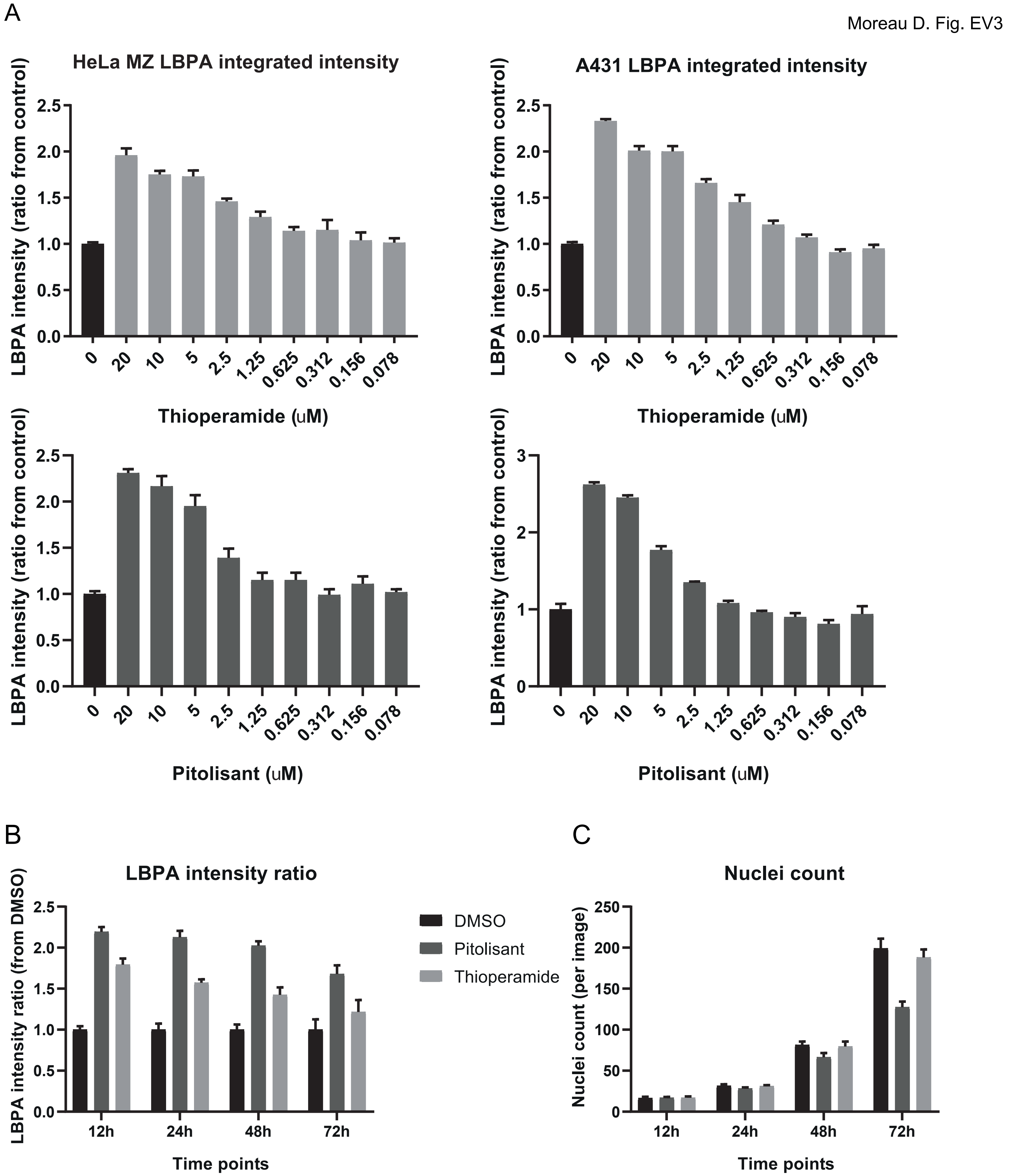

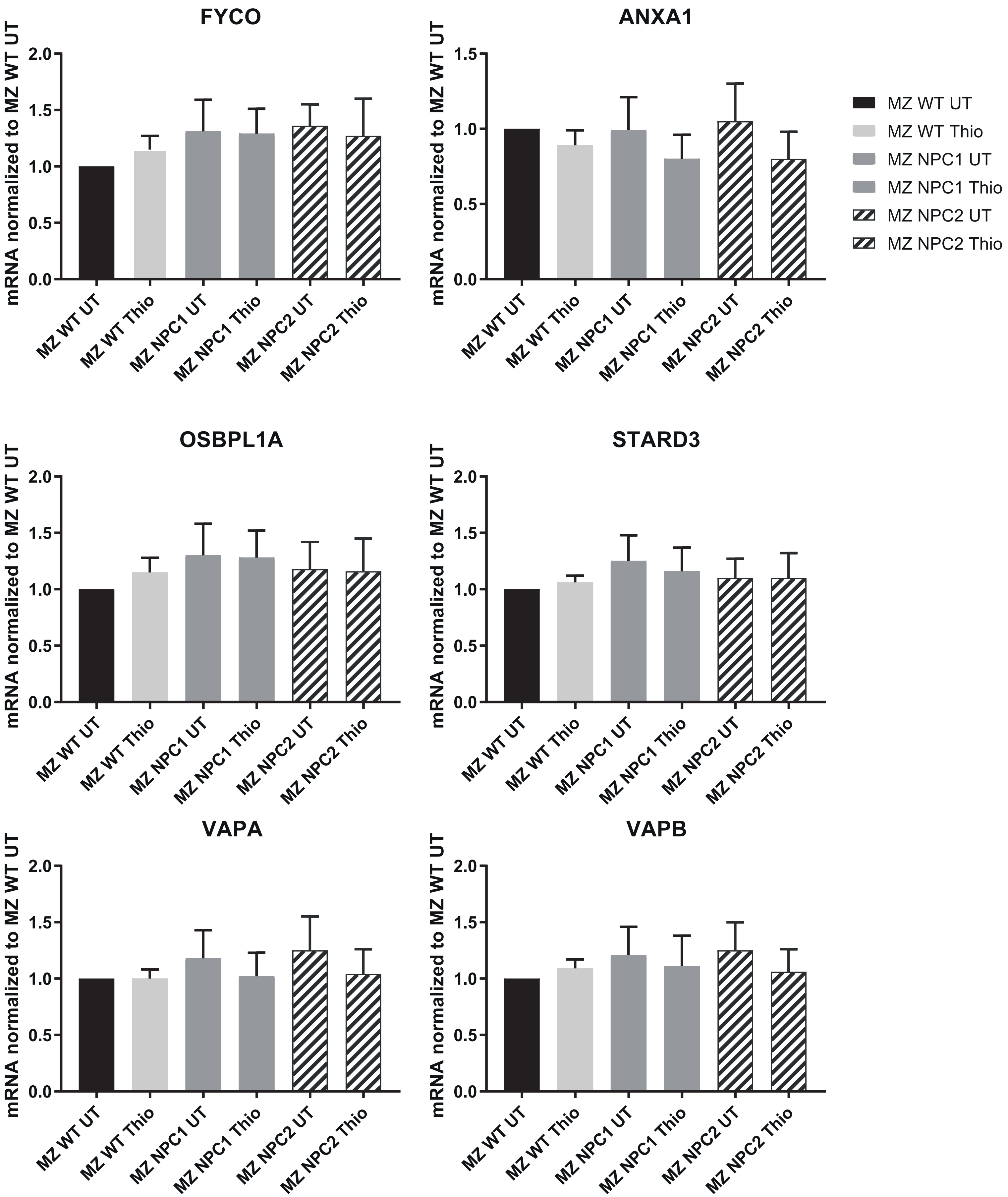

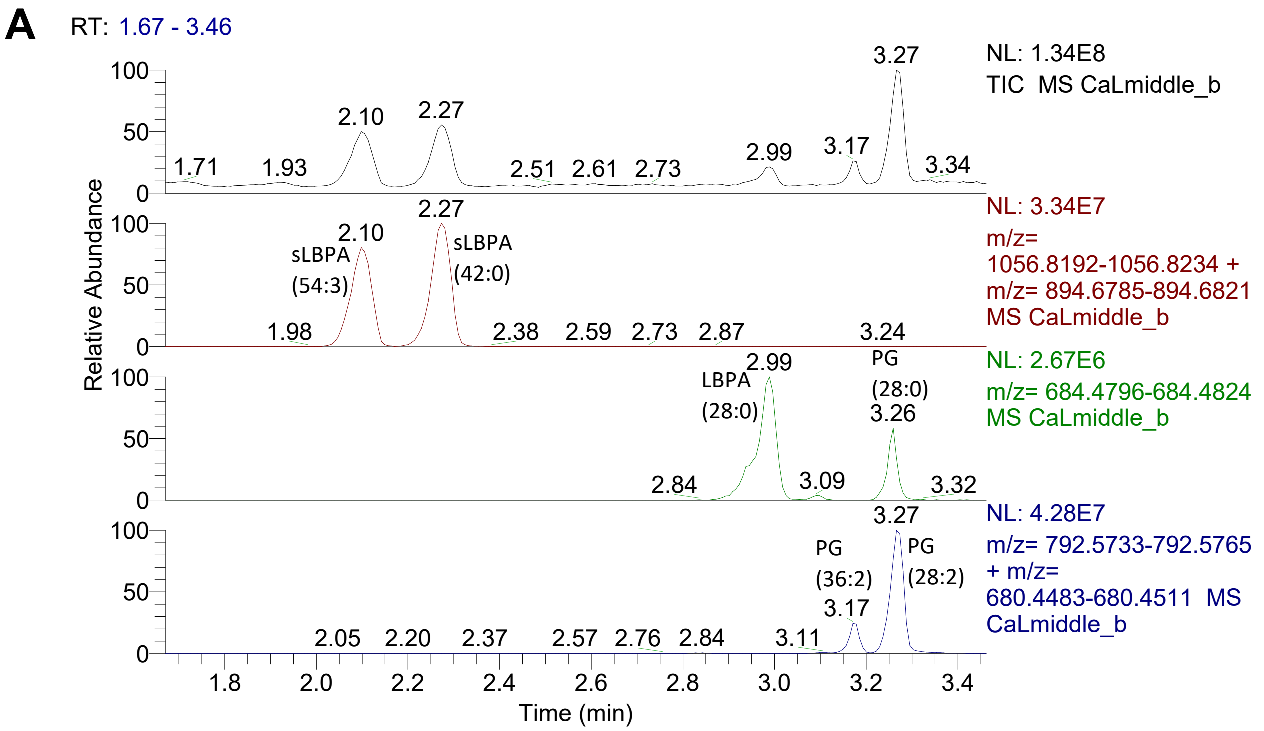

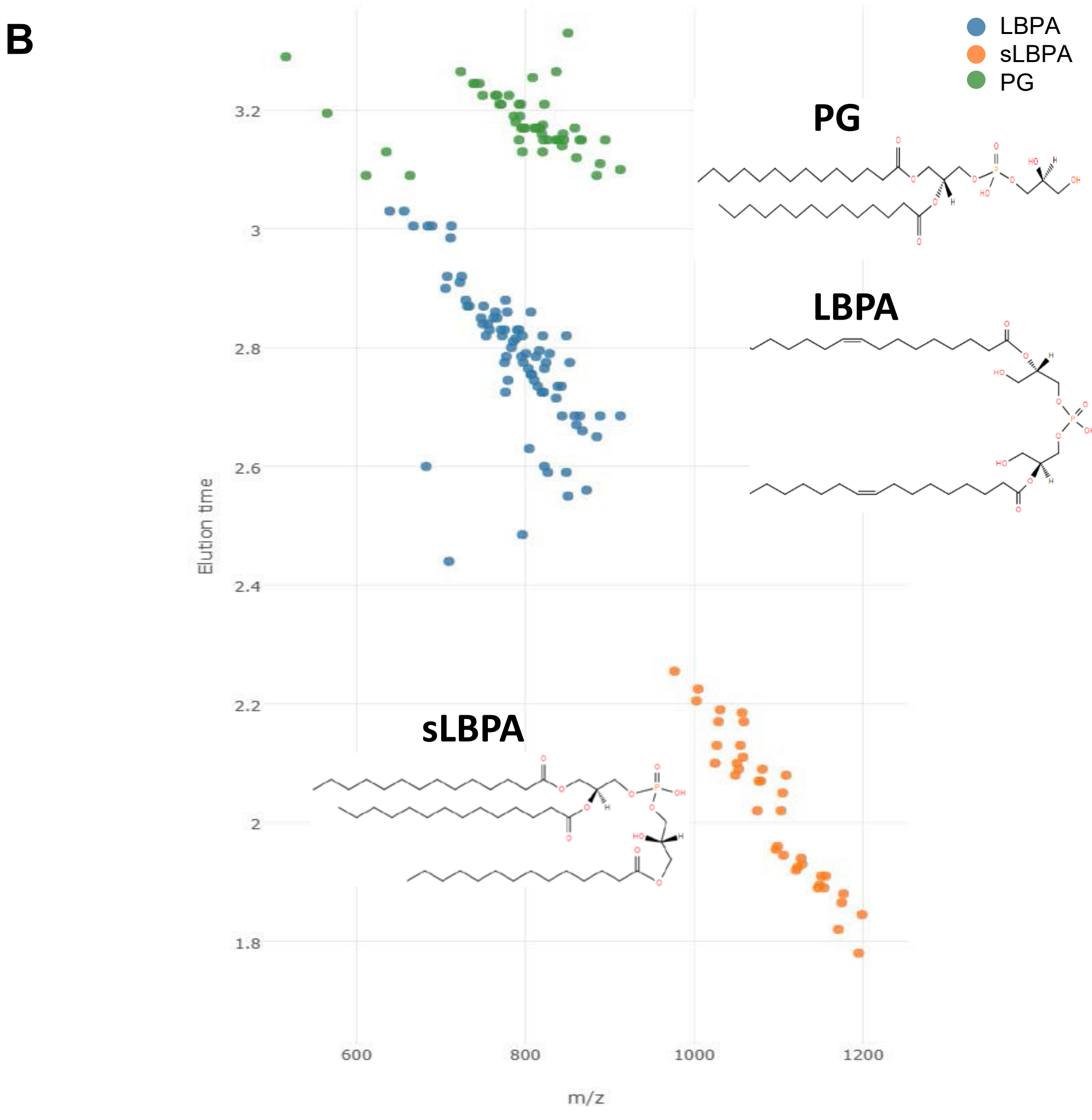

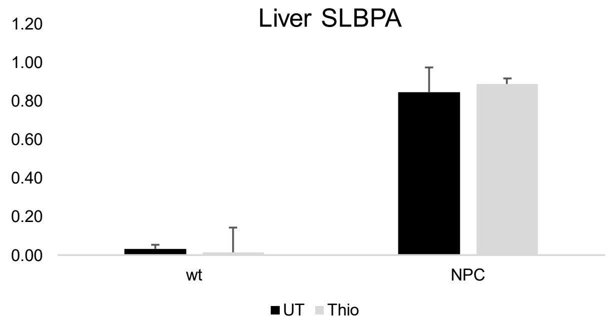

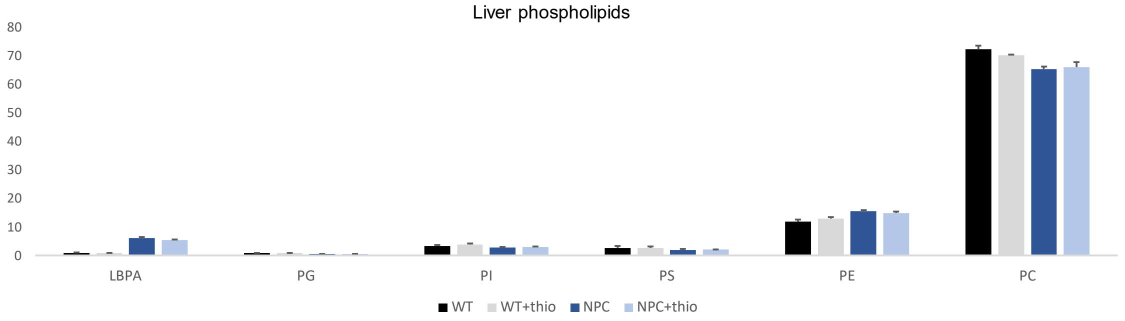

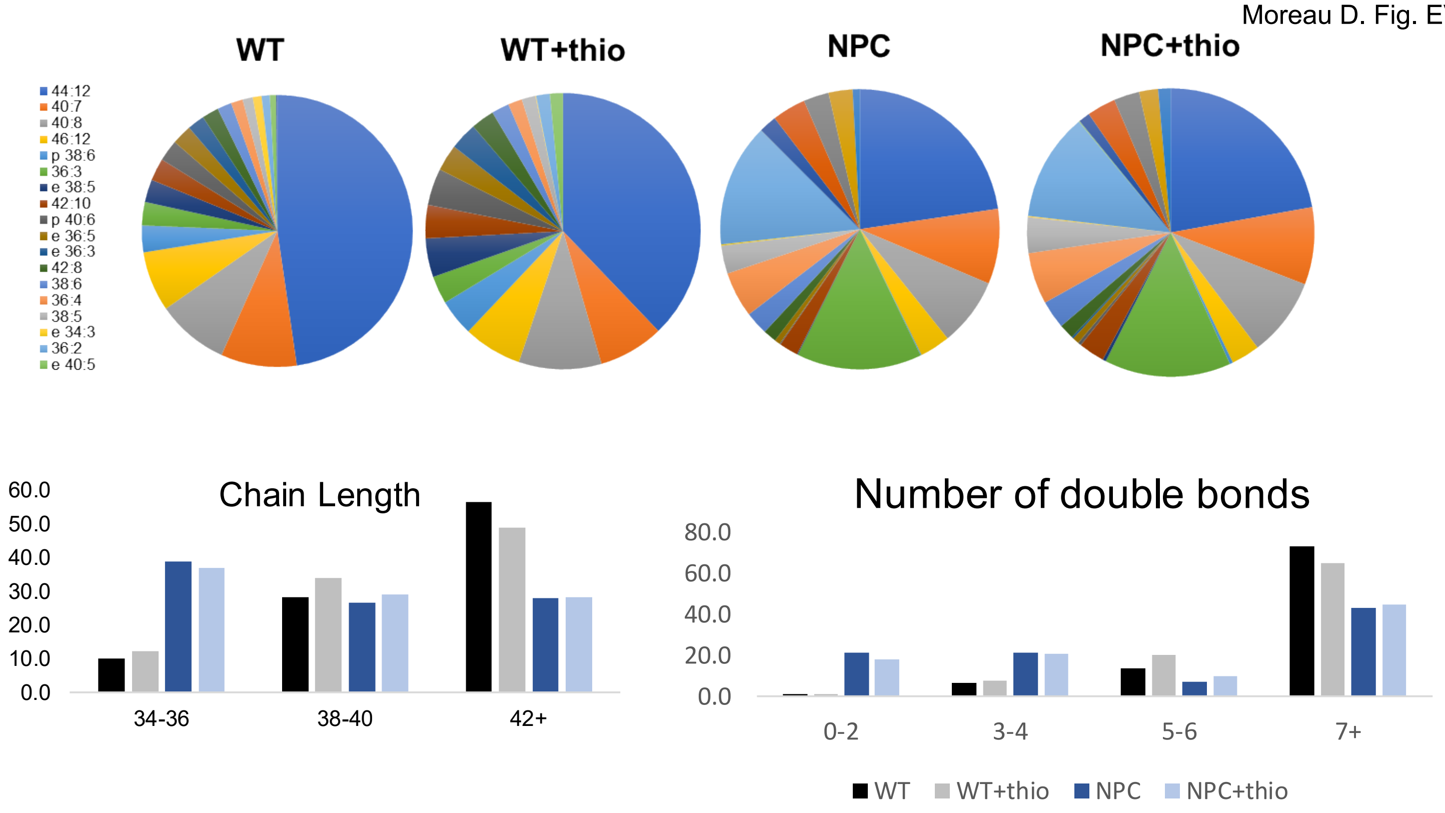

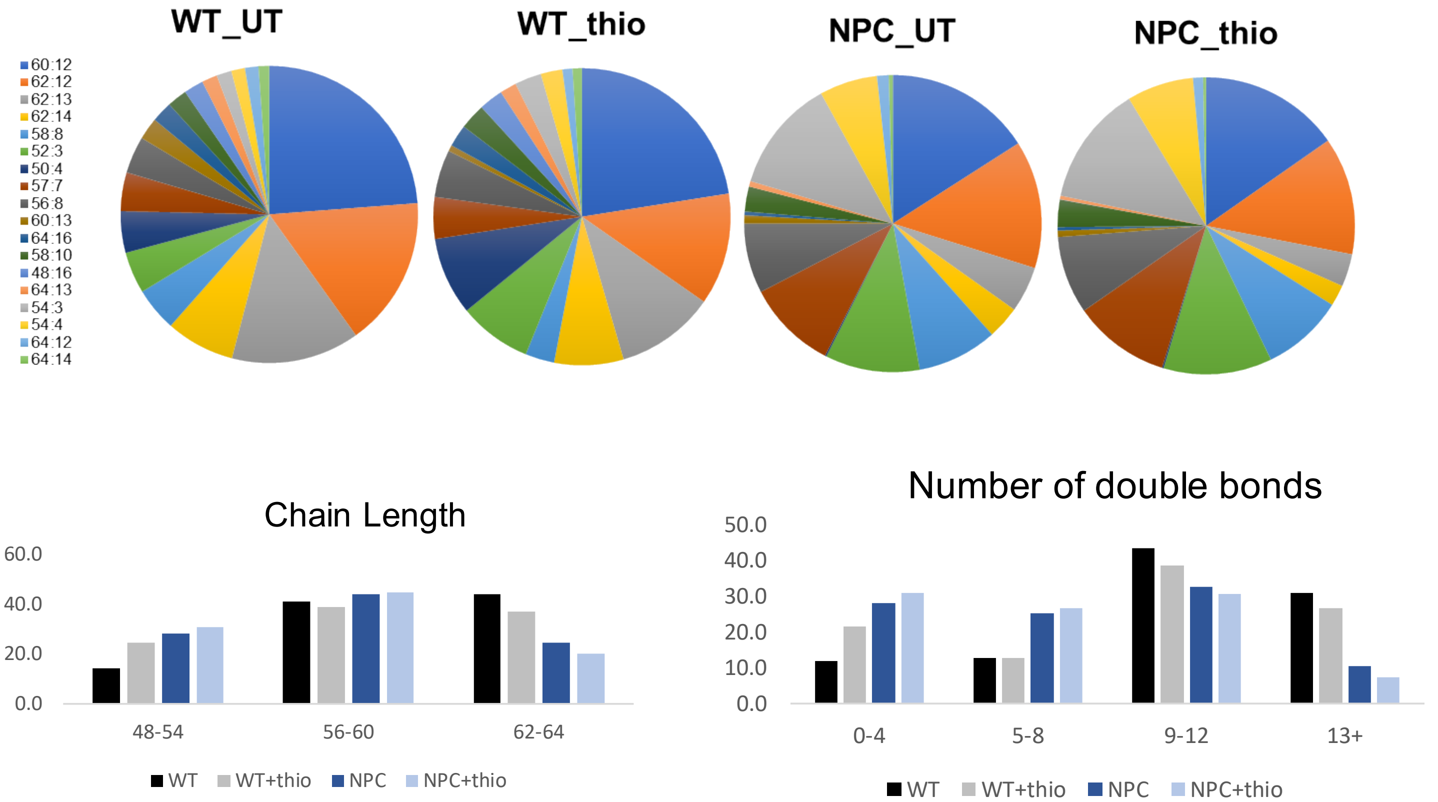

Moreau D. Fig. EV8

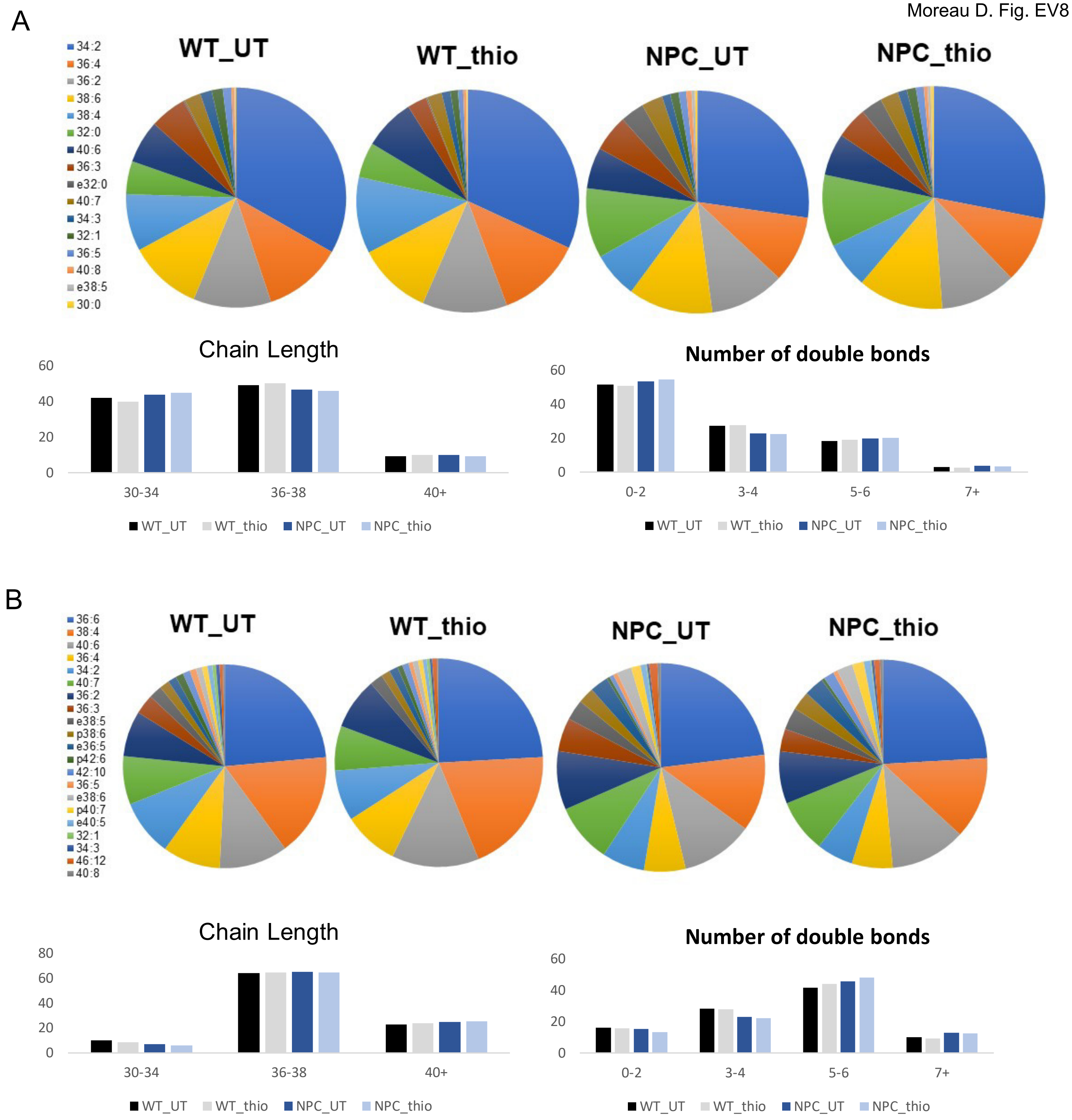

Moreau D. Fig. EV9

A

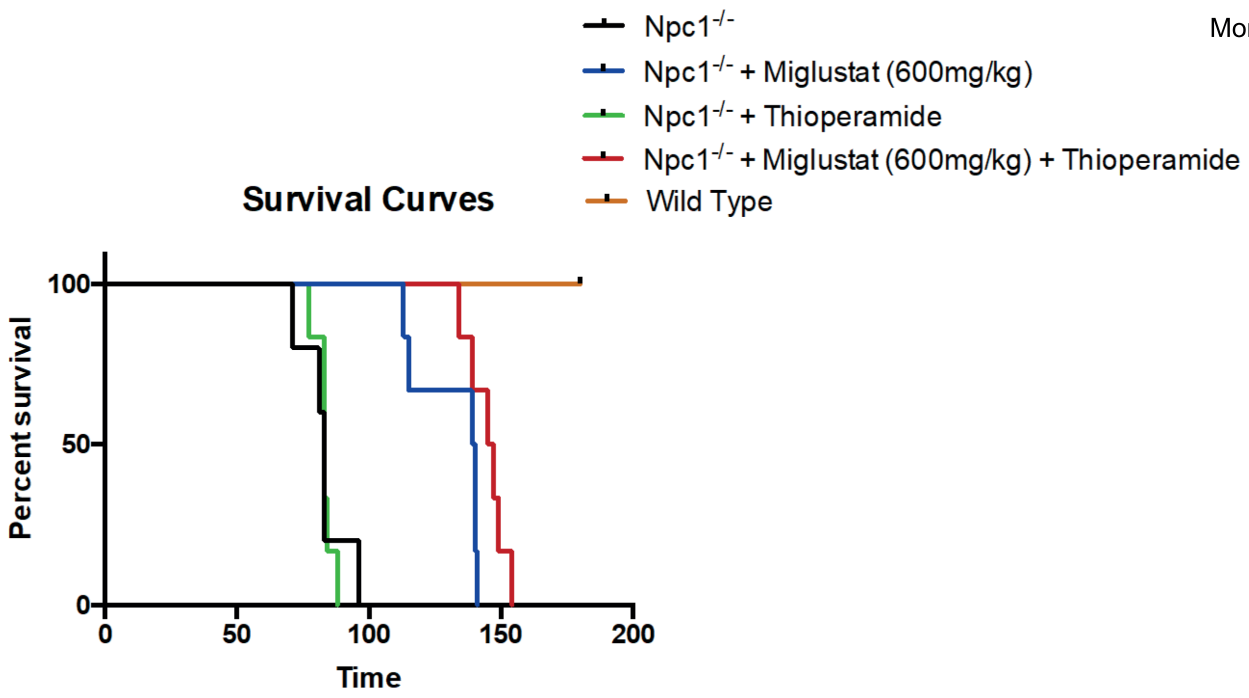


B

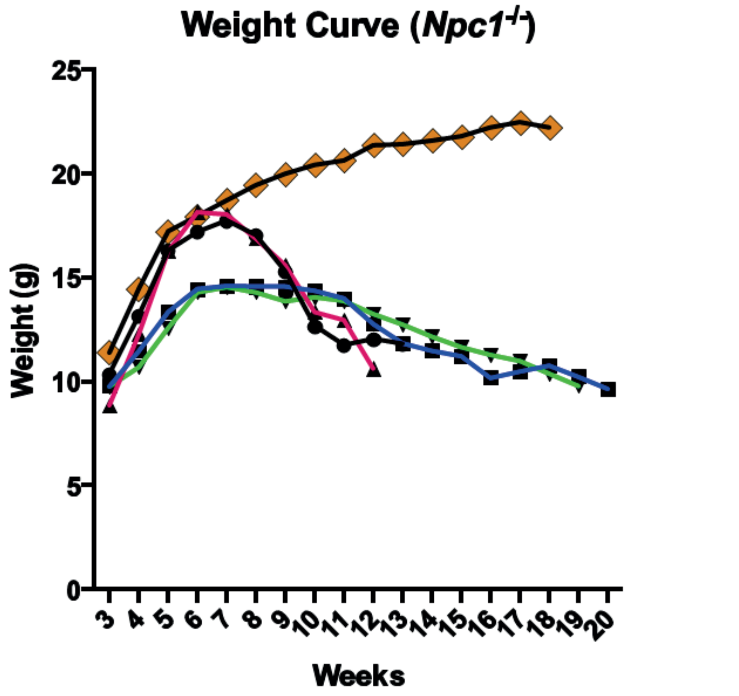


C

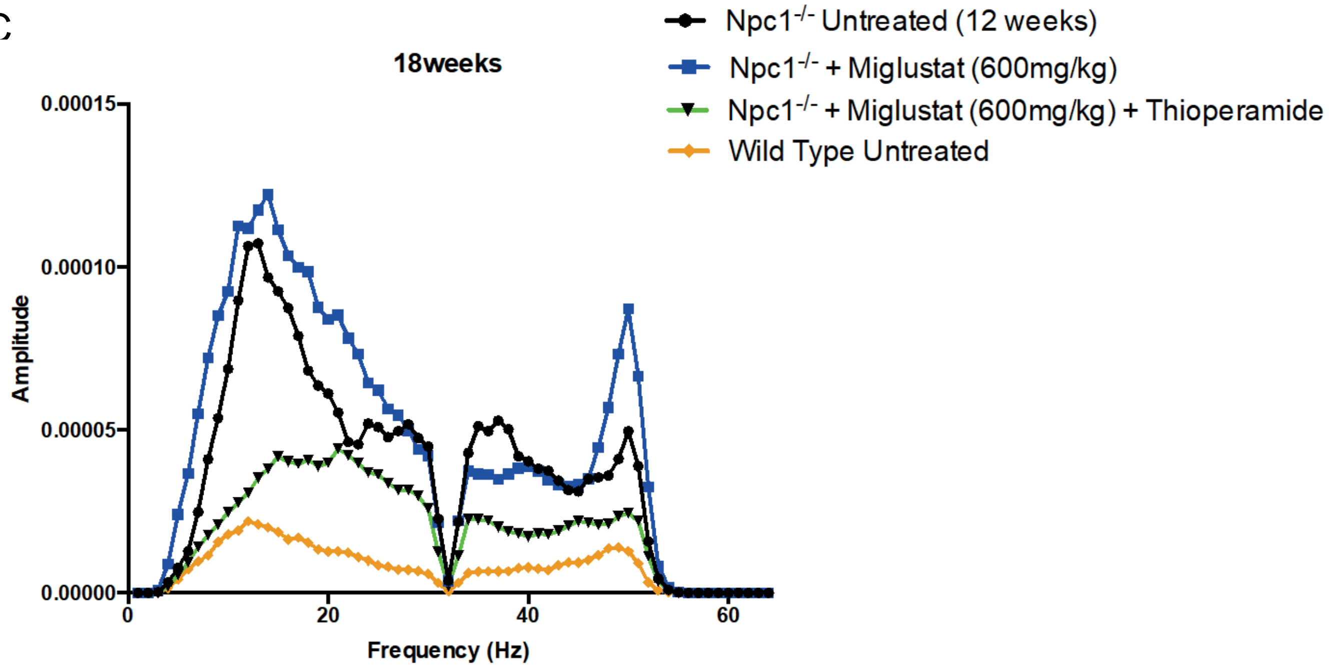

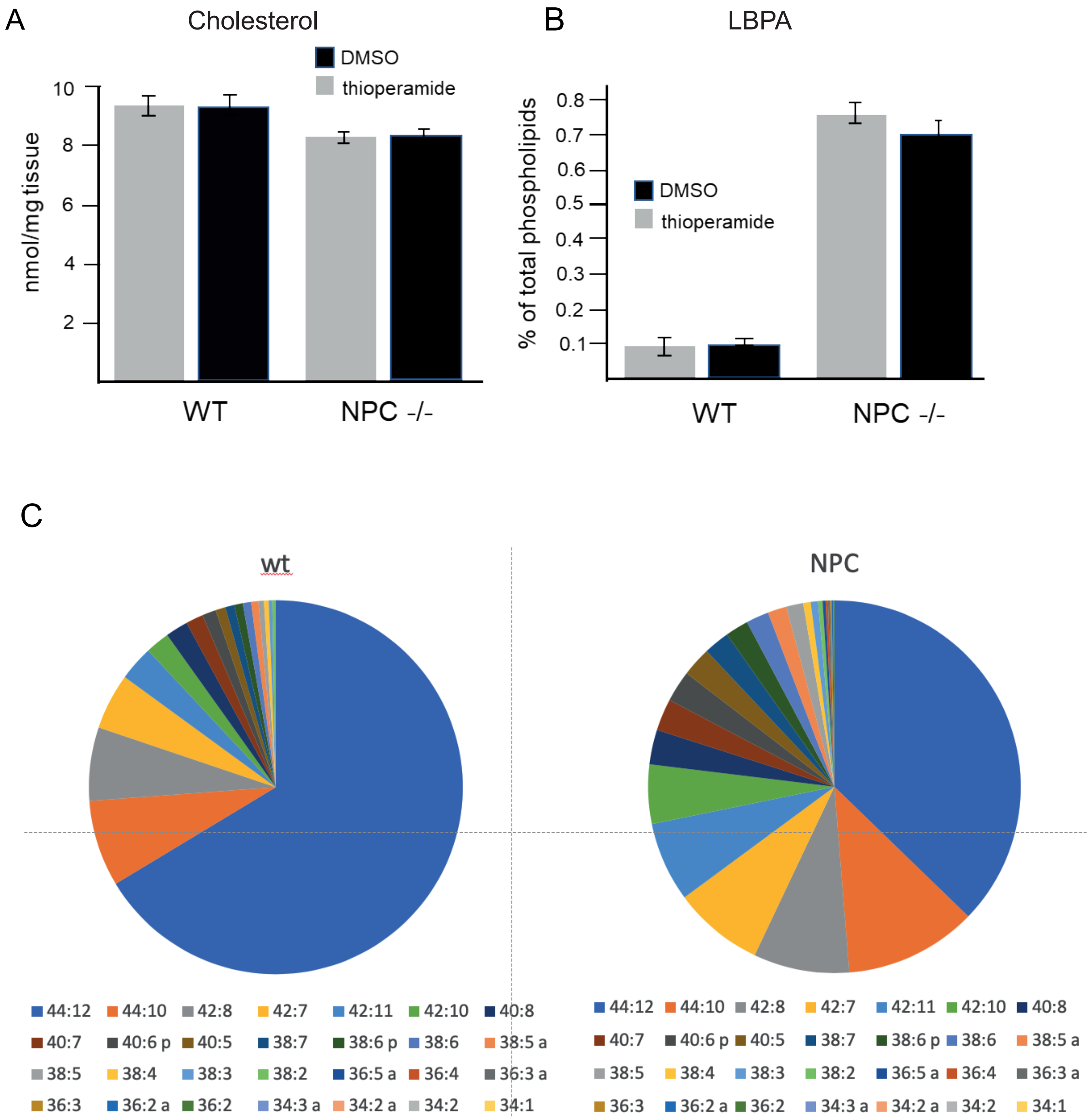
